# Nociceptin Orphanin F/Q Pathways are Dysregulated by Stress and Modulate Reward Learning and Motivation Across Species

**DOI:** 10.1101/2025.05.05.652258

**Authors:** Diego A. Pizzagalli, Michael T. Treadway, Brian D. Kangas, Benedetto Romoli, Jocelyn Breton, Michael R. Bruchas, Ann M. Graybiel, Emily Hueske, Nandkishore Prakash, Andre Der-Avakian, Sabina Berretta, Kevin G. Bath, Davide Dulcis

## Abstract

Nociceptin orphanin F/Q has been implicated in stress-related depressive phenotypes. Specifically, exposure to chronic stressors upregulates nociceptin receptors (NOPR), whereas NOPR antagonism has anti-depressant/anti-anhedonic effects. The mechanisms underlying these effects remain, however, unclear. Here, we investigated the role of NOPR in depressive phenotypes alongside potentially prohedonic effects of NOPR antagonism across species. In Study 1, we evaluated whether exposure to early-life adversity (ELA) upregulated ventral tegmental area (VTA) and striatal prepronociceptin (*Pnoc*) gene expression in adult mice. In Study 2, we tested whether chronic social defeat altered *Pnoc* gene expression in reward-related regions. To establish whether direct NOPR modulation is implicated in reward-related behaviors, in Study 3, we assessed whether NOPR antagonism alters reward learning in rats. Finally, in Study 4, we tested whether NOPR antagonism boosts motivation among depressed humans. ELA induced anhedonic behavior and increased *Pnoc* expression in the VTA; in females (but not males), ELA increased *Pnoc* expression in the dorsal striatum (Study 1). Furthermore, chronic stress reduced *Pnoc*-expressing cells in the VTA, dorsal striatum and prefrontal cortex and susceptible rats showed reduced VTA NOPR gene (*Oprl1)-*expressing cells (Study 2). In a behavioral assay, a single 30-mg dose of a NOPR antagonist (BTRX-246040) boosted reward learning in rats (Study 3). Finally, in depressed humans, relative to placebo, 8-week treatment with BTRX-246040 increased incentive motivation (Study 4). Collectively, our findings indicate that chronic stressors alter *Pnoc* and mRNA levels of *Pnoc*-expressing cells in a sex-selective and region-specific manner impacting reward structures, and that NOPR antagonism shows anti-anhedonic properties.

## Introduction

Despite decades of research, the pathophysiology of major depressive disorder (MDD) remains incompletely understood, and treatment advances have been limited. Substantial preclinical evidence implicates nociceptin/orphanin FQ peptide (N/OFQ) and its endogenous receptor (NOPR) in the emergence and maintenance of depressive and anhedonic phenotypes as well as a promising antidepressant target [1–3]. Several findings are consistent with these assumptions. First, N/OFQ and NOPR are known to be critically involved in regulating stress, reward processing, and pain [1,2] – domains that are impaired in MDD [4–6]. Second, various stressors known to induce depression-like behaviors in rodents have been found to upregulate N/OFQ [7]. For example, rats exposed to a 3-day of social defeat were characterized by increased N/OFQ peptide-encoding *Pnoc* mRNA levels in the striatum as well as reduced reward learning. In addition, N/OFQ peptide and NOPR gene-encoding *Oprl1* mRNA levels in the anterior cingulate cortex (ACC), ventral tegmental area (VTA) and striatum were negatively correlated with amount of reward learning [8]. Similarly, chronic social defeat as well as administration of a NOPR agonist both induced depressive phenotypes in mice, including reduced social interaction and exploration, and both increased pro-inflammatory cytokines (Camara & Brandao, 2025). Conversely, NOPR knockout mice have been characterized by antidepressant-like phenotypes in behavioral despair assays [9]. Third, administration of NOPR antagonists produced antidepressant-like effects in mice [9–11], prevented the emergence of helplessness during inescapable stress exposure in mice [12,13], and reversed stress-induced anhedonic phenotypes (decreased sucrose preference) in rats [14]. Of note, administration of BTRX-246040 (formerly known as LY2940094) – a NOPR antagonist with high affinity, potency and selectivity for the NOPR over other opioid receptors [15,16] – before stress exposure prevented the development of depressive phenotypes (helplessness behavior) [17]. These findings extend additional preclinical evidence in rodents highlighting that BTRX-246040 exerts robust antidepressant and anti-anhedonic effects [18–20]. Finally, initial data indicate that chronic administration of a NOPR antagonist in patients with MDD has antidepressant effects [21], although this effect was not replicated in an independent Phase 2 clinical trial (https://clinicaltrials.gov/study/NCT03193398).

Additional evidence for a key role of the nociceptin system in the pathophysiology of MDD stems from anatomical studies highlighting NOPR expression distributed along brain circuitry that are abnormal in MDD, including the prefrontal cortex (PFC), hippocampus, amygdala, dorsal raphe nucleus, VTA, and striatum. Thus, autoradiography studies in rats have demonstrated a high concentration of NOPR in the frontal cortex and limbic structures (including the hippocampus and amygdala) [22,23]. Fitting these data from rodents, human positron emission tomography and post-mortem autoradiography studies have shown high concentrations of NOPR in the PFC and striatum [24,25]. NOPR have been localized in serotonergic, noradrenergic, and dopaminergic neurons, including in the raphe complex, locus coeruleus, VTA and substantia nigra [22,26]. Critically, both *in vitro* and *in vivo* studies across several brain regions have clarified that activation of NOPR by N/OFQ or by an agonist produces a generalized inhibition of neurotransmitter release, including dopamine and serotonin but also glutamate and GABA [3].

Particularly relevant to the current studies, studies have demonstrated high expression of N/OFQ and NOPR in brain regions of the mesocorticolimbic dopaminergic pathways, which are critically implicated in reward learning and motivated behaviors [3,27]. In this context, N/OFQ has been found to inhibit DA release in the VTA, nucleus accumbens and caudate nucleus [18–20], and contributes to dopamine cellular loss in the striatum [31]. Conversely, NOPR antagonists enhanced DA transmission [32,33]. In light of the key role of dopamine in reward learning and motivation [34,35], relationships between N/OFQ and dopamine might be particularly important in the context of anhedonic phenotypes [36]. Consistent with this assumption, N/OFQ neuron activation in the pnVTA was found to constrain motivation for reward [37] and single dose of a NOPR antagonist potentiated electrophysiological markers linked to positive prediction errors implicated in reward learning [38]. Collectively, these data suggest that N/OFQ upregulation might constitute a key pathophysiological mechanism linked to depression vulnerability and that NOPR antagonism could represent an exciting and relatively novel antidepressant treatment option, especially targeting anhedonic and/or motivational deficits [1,2].

In spite of this robust preclinical evidence of a critical role for N/OFQ and NOPR in anhedonia and depression-related behaviors, several unanswered questions remain. First, it is unclear whether N/OFQ and NOPRs are affected by early-life adversity, which is a well-established and potent risk factor for MDD later in life [39] and likely increases risk for depression via disruption of reward-related pathways and reward learning (among other mechanisms) [40–42]. Intriguingly, we recently reported that, during early development, *Pnoc* is strongly expressed in a population of dopamine-projecting striatal neurons (striosomes) shown to wield powerful control over nigral dopamine neurons [43–45]. Second, knowledge gaps remain with respect to co-localization of neurons expressing *prepronociceptin* (*Pnoc*) – the gene encoding pre-pro-N/OFQ, the precursor polypeptide of the synaptically-acting neuropeptide neuromodulator, N/OFQ, with highest affinity for activating NOPR – and its receptor across brain regions relevant to stress-related depressive phenotypes. Third, no study has evaluated whether NOPR antagonism boosts hedonic behavior (reward learning) in an assay modeled after tasks used in humans with MDD and anhedonia [46,47]. Finally, there are no prior reports evaluating whether sustained treatment with a NOPR antagonism can boost motivation (willingness to exert efforts for rewards) in individuals with MDD. Here, we present findings from four complementary studies, across species and domains, to answer these key questions in the field by establishing the sensitivity of the N/OFQ-NOPR system to stress, and whether perturbing this system could have beneficial effects on motivation in depressed human individuals.

Thus, in Study 1, we applied real-time quantitative polymerase chain reaction (RT-qPCR) to test the hypothesis that exposure to early-life adversity (limited bed nesting (LBN)) would upregulate striatal *Pnoc* gene expression later in life in mice. In Study 2, we used RNA-scope in-situ hybridization to test the hypothesis that chronic social defeat would alter *Pnoc* gene expression in reward-related regions. To this end, we quantified mRNA levels by *in situ* hybridization (ISH) using RNAscope probes labeling *Pnoc* (*Pnoc*-ISH), NOPR mRNA (Oprl1-ISH), *c-Fos*, and neurotransmitter markers for dopamine (tyrosine hydroxylase mRNA, TH-ISH), glutamate (vesicular glutamate transporter1 mRNA, VGLUT1-ISH), and GABA (vesicular GABA transporter mRNA, VGAT-ISH) in mice exposed to 21-day chronic social defeat; analyses focused on the VTA, dorsal striatum and PFC and intracranial self-stimulation (ICSS) was used to categorize susceptible vs. resilient rats. In Study 3, we tested the hypothesis that, relative to vehicle, high doses of a NOPR antagonist (BTRX-246040) would boost reward learning among control (unstressed) rats tested with the rodent version of the Probabilistic Reward Task (PRT). Finally, in Study 4, we evaluated the hypothesis that 8-week treatment with the same NOPR antagonist would boost willingness to expend efforts for rewards in individuals with MDD.

### Study 1: Adverse, Unpredictable Early-life Experiences Upregulates Dorsal (But Not Ventral) Striatal *Pnoc* Gene Expression in Female (but Not Male) Mice

#### Mice

Male and female C57BL/6N mice were bred in house, had *ad libitum* access to food and water, and were housed on a 12 h:12 h light:dark cycle. All animal procedures were approved by the New York State Psychiatric Institute (NYSPI) Institutional Animal Care and Use Committee (IACUC) and were consistent with the guide for the care and use of animals in research.

#### Limited bedding and nesting (LBN)

Four days following birth of a litter, the dam and pups were transferred from their standard home cage to a home cage with a wire mesh floor and a 2 × 4 cm cotton nestlet as their only source of bedding (Bath et al., 2016). Mice continued to have *ad libitum* access to food and water. Dam and litters remained in these modified housing conditions for seven days, and were then returned to standard housing, containing cob bedding and a 4 × 4 cm nestlet. Control mice were left undisturbed throughout these procedures. Litters were composed of both male and female pups and litters ranged in size from 5 to 8 pups per litter. All pups were weaned and sex segregated at 21 days of age.

#### Real-time quantitative polymerase chain reaction (RT-qPCR)

For gene expression studies, adult male and female mice reared under either control or LBN conditions. After rapid decapitation, the brains were collected, flash frozen, blocked, and tissue punches were collected from the dorsal and ventral striatum. Tissue collection was completed from at least three different litters per condition to eliminate the possibility of cohort effects on measures of gene expression (n = 8-12 animals per group). Tissue punches were placed in RNAzol (Molecular Research Center, Cincinnati, OH), homogenized by brief sonication and stored at −80 °C until further processing. RNA isolation was in accordance with the manufacturers protocol. First strand cDNA synthesis was in accordance with New England Biolabs MmULV protocols (NEB, Ipswitch, MA). We used pre-designed and pre-validated Taqman assays from Applied Biosystems (Life Technologies, Norwalk, CT) run in multiplex with housekeeping gene (18S). For each plate and assay, gene expression was calculated relative to a standard curve included on each plate. We used a CFX384 RT-qPCR system (Biorad, Hercules, CA) and associated software for all gene expression profiling.

#### Assessment of anhedonic phenotype (sucrose preference)

Control and LBN animals were housed in clean cages and given 48 hours to habituate to the two-bottle paradigm. Two metered bottles (Animal Care Systems, Part: C79125) were placed in the cage; one bottle contained a 1% sucrose solution and the other contained plain, unsweetened drinking water. Following the two days of habituation, mice were tested across a 72-hour test period. Animals were given free access to both bottles throughout all days of habituation and testing. Animals were not food or water deprived. To prevent possible side-preferences, the position of the bottles in the cage were switched every 24 hours. Bottles were weighed at the same time each day, and the intake of water and 1% sucrose, as well as sucrose preference (percent sucrose consumed/total intake) was measured. Sucrose preference was averaged across the three days of the test period.

#### Statistics

All data are presented as the mean ± the standard error of the mean. One outlier datapoint (one LBN male mouse) was removed from the analysis from the *Pnoc* gene expression analysis because its data were outside two standard deviations from the mean. Two-way analysis of variance (ANOVA) was used to test for effects of *Rearing Condition* (LBN, control) and *Sex*. Statistically significant main effects were followed up with Sidak post-hoc comparisons. An alpha of 0.05 was used for all analyses. All statistics were performed using GraphPad Prism version 9.2 (GraphPad Software, San Diego, California USA).

## Results

### Pnoc gene expression

Tissue from the VTA as well as dorsal and ventral striatum was collected from adult male and female mice reared in either control or LBN conditions. RT-qPCR was used to assess changes in striatal *Pnoc* gene expression across groups. In the dorsal striatum, a two-way ANOVA identified a statistically significant *Rearing Condition* x *Sex* interaction (*F*(1, 37) = 11.19, *p* = 0.0019, 1^2^ = 0.23; Figure 1A), but no main effect of *Rearing Condition* or *Sex* (both Fs < 0.07, *p*s > 0.75). Subsequent post-hoc tests identified a significant increase in *Pnoc* expression in LBN females relative to controls (*p* = 0.022; *d* = 1.11) and a trend for males (*p* = 0.08; *d* = 1.01) (Figure 1A). In the ventral striatum, there was a main effect of *Sex* (*F*(1,36) = 5.49, *p* = 0.025, 1^2^ = 0.13) with males showing elevated *Pnoc* compared to females (males: 1.65± 0.34; females: 0.91±0.09). However, there was no effect of *Rearing Condition* and no significant interaction (both Fs < 1.20, *p*s > 0.30; Figure 1B). In the VTA, a main effect of *Rearing Condition* emerged, driven by overall higher *Pnoc* expression in LBN than control mice (*F*(1,39) = 4.66, *p* = 0.037, 1^2^ = 0.104; Figure 1C).

**Figure 1:**
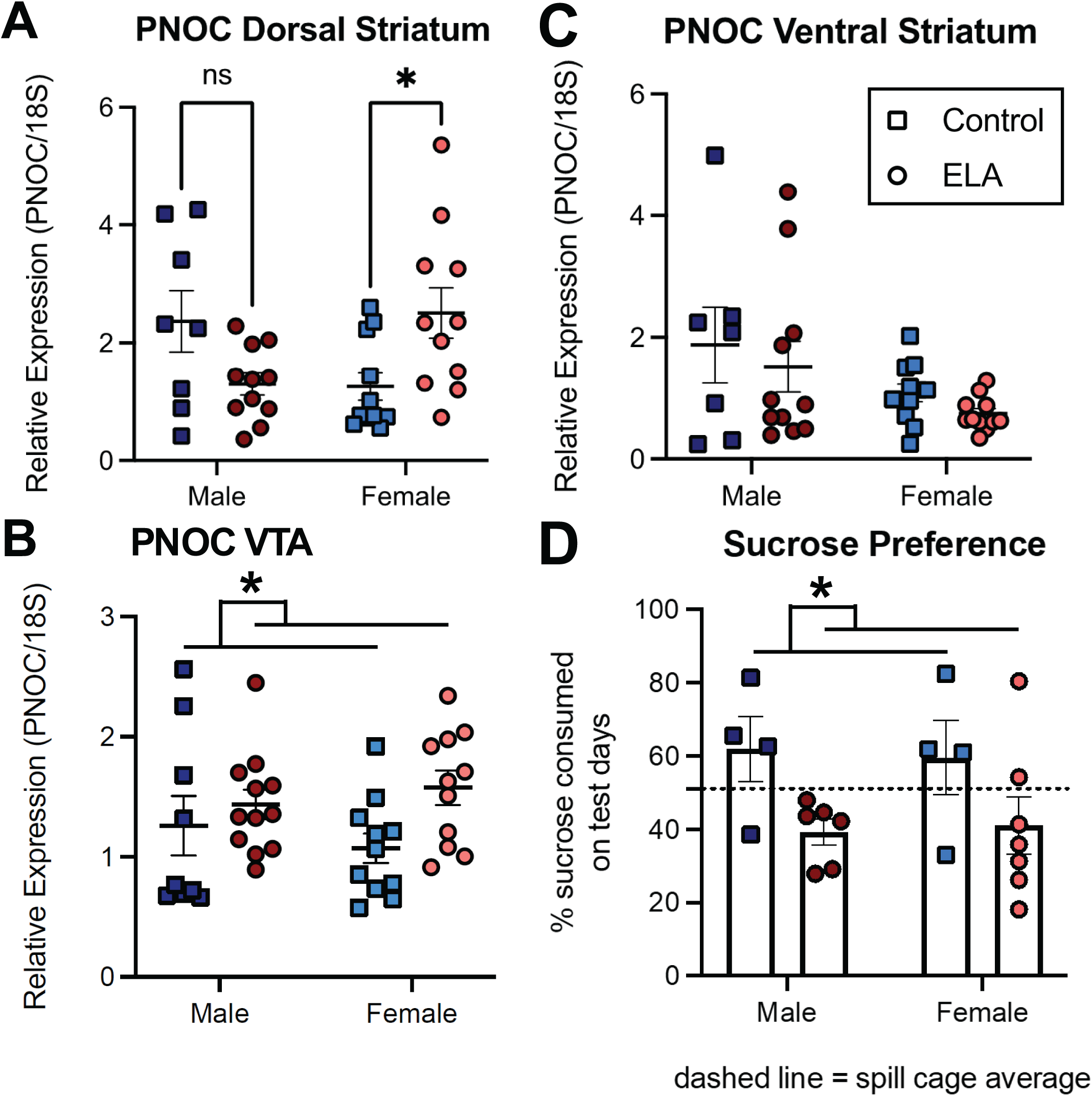
**(A)** Relative to female controls, LBN females were characterized by increased *Pnoc* expression in the dorsal striatum. This effect was not seen in LBN males (significant *Sex* x *Rearing Condition* interaction). **(B)** Relative to control mice, LBN mice were characterized by increased *Pnoc* expression in the VTA. **(C)** Relative to control mice, LBN mice did not differ in their *Pnoc* expression in the VTA. **(D)** In both male and female mice, the LBN procedure induced anhedonic behavior (reduced sucrose preference).

### Anhedonic behavior (sucrose preference)

Across the three days of testing, animals reared under LBN conditions demonstrated reduced sucrose preference relative to control animals, as evidenced by a statistically significant main effect of *Rearing Condition* (*F*(1,17) = 6.98, *p* = 0.017, 1^2^ = 0.29; Figure 1D). There was no main effect of *Sex* and no significant interaction between *Rearing Condition* and *Sex* (both Fs < 0.08, *p*s > 0.75). LBN-induced anhedonia was not sex-specific, as both males and females reared under LBN conditions displayed lower sucrose preferences, on average, compared to their control counterparts (control males: 62.0±8.85; LBN males: 39.23± 3.48; control females: 59.6±10.12; LBN females: 41.06± 7.86).

### Study 1 Interim Discussion

Results from Study 1 show that adult animals reared under LBN conditions display an anhedonic phenotype (decreased sucrose preference), consistent with prior studies using this assay of early-life adversity in both rats and mice [48,49]. Of note, in Study 1, such anhedonic behavior emerged in both male and female mice, whereas some prior studies have demonstrated anhedonia for natural rewards (e.g. palatable food and social play) and drugs (e.g. cocaine) in male but not female rats [50–52]. In addition, a recent study reported that LBN led to enduringly blunted reward learning [53], which was driven by reduced sensitivity to rewards [54]. Consistent with our hypotheses, relative to control rearing, rearing under LBN conditions was associated with overall higher *Pnoc* expression in the VTA, whereas in the dorsal (but not ventral) striatum LBN upregulated *Pnoc* in female (but not male) mice, highlighting sex- and region-specific selectivity. Although these findings establish that a model of early adversity affects *Pnoc* expression in the VTA and dorsal striatum among adult mice, it is unclear if similar effects (1) emerge in a different species (rats), in adult animals exposed to stress, and in additional regions critically implicated in the pathophysiology of MDD, (2) can be elicited by a different chronic stressor, (3) are manifested in a different anhedonic phenotype, and (4) involve additional neurotransmitters. Thus, in Study 2, we used RNAscope *in-situ* hybridization to quantify neurons expressing *Pnoc* and its receptor *Oprl1* in multiple brain regions of a model of social defeat-induced stress, and probed anhedonic behavior using ICSS. In addition, we evaluated whether putative abnormalities are specific to vulnerable vs. resilient animals and affect GABA and glutamate, in addition to dopamine.

### Study 2: Social Defeat Affects Transcription of Nociceptin Peptide and *Oprl1* Receptor in Reward-Processing Brain Areas

In Study 2, rats underwent 21 days of social defeat while being tested for ICSS to measure anhedonia. Critically, by the end of the chronic stress protocol, approximately half of the stressed animals showed a susceptible phenotype (increased anhedonia), whereas the remaining animals showed a resilient phenotype (they showed anhedonia levels comparable to controls).

#### Methods

Adult male Wistar rats (N = 12) were subjected to 21 days of social defeat and anhedonia was objectively assessed using the ICSS procedure. As shown in Supplemental Figure 1, the stress-exposed animals clustered into 50% vs. 50% susceptible vs. resilient rats based on their ICSS reward threshold (unbiased k-means method). Brain sections through the levels of the VTA, dorsal striatum and PFC were processed for RNA-scope in-situ hybridization to detect mRNA levels of *Pnoc* (PNOC-ISH), *Oprl1* (Oprl1-ISH), *c-Fos*, and neurotransmitter markers for dopamine (*TH*-ISH), glutamate (*VGLUT1*-ISH), and GABA (*VGAT*-ISH).

### Study 2 Results

Quantification of in-situ hybridization-reactive cells in these stress- and reward-related brain regions revealed that stress induced a significant reduction of *Pnoc*-expressing cells in VTA, dorsal striatum and PFC (Figure 2), while no significant changes were detected in the number of *Oprl1*-expressing cells (Figure 3A-C). When stressed animals were clustered into susceptible and resilient groups (Supplemental Figure 1), analyses showed that *Pnoc* transcriptional reduction in all three brain regions was driven by the resilient group; specifically, susceptible animals continued to have higher levels of *Pnoc* than resilient rats, which is consistent with the higher *Pnoc* in LBN mice emerging from Study 1. The same data cluster analysis on *Oprl1* expression showed that it was differentially regulated in specific brain under different conditions (Figure 3D). Susceptible rats had significantly fewer VTA *Oprl1*-expressing cells than resilient rats; resilient rats, by contrast, had significantly fewer dorsal striatum *Oprl1*-expressing cells than the susceptible rats.

**Figure 2:**
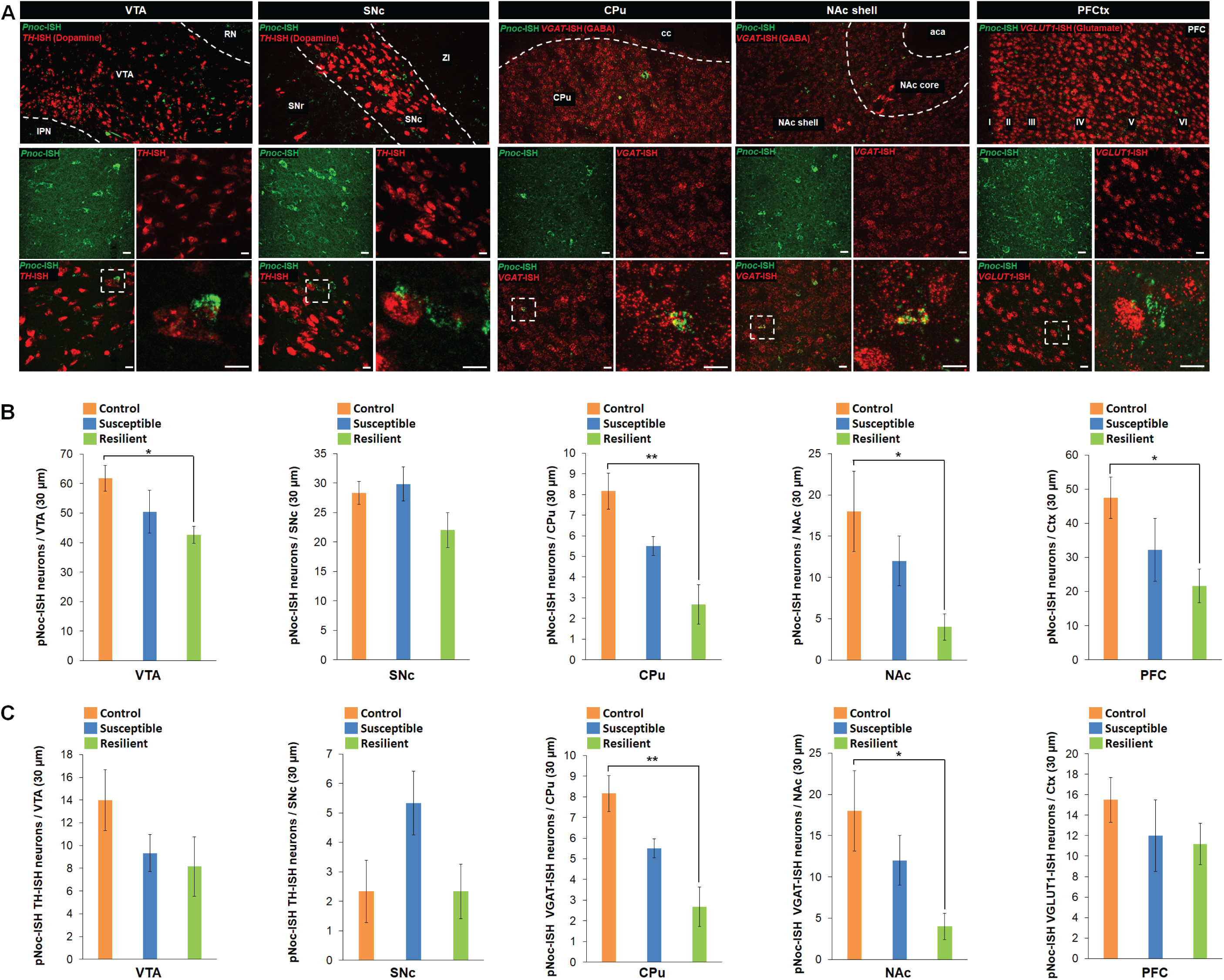
**(A)** Confocal images of *Pnoc* (green) RNAscope in-situ hybridization (PNOC-ISH) in Ventral Tegmental Area (VTA), Substantia Nigra pars compacta (SNc), Caudate-Putamen (CPu), Nucleus Accumbens shell (NAc) and Prefrontal Cortex (PFC). The main NT-expressing cell type for each brain region was counterstained in red: Tyrosine hydroxylase (Th-ISH) for VTA and SNc; Vescicular GABA transporter (VGAT-ISH) for CPu and NAc; Vescicular glutamate transporter 1 (VGLUT1-ISH) for PFC. **(B)** Quantification of neurons expressing *Pnoc* transcript in Control (N=6), susceptible (N=6) and resilient (N=6) rats for each brain region. ICSS-based clustering into susceptible and resilient groups revealed that the decrease in Pnoc-ISH+ neurons is found mostly in resilient rats. Tukey’s multiple comparisons, * p<0.05, ** p < 0.01. **(C)** Quantification of NT-expressing *Pnoc*+ neurons (TH for VTA, SNc; VGAT in CPu, NAc; VGLUT1 in PFC) in Control (N=6), susceptible (N=6) and resilient (N=6) rats for each brain region. The decrease of *Pnoc*+ neurons did not occur within these NT-expressing neuronal pools in VTA and PFC. Tukey’s multiple comparisons, * p<0.05, ** p < 0.01.

**Figure 3:**
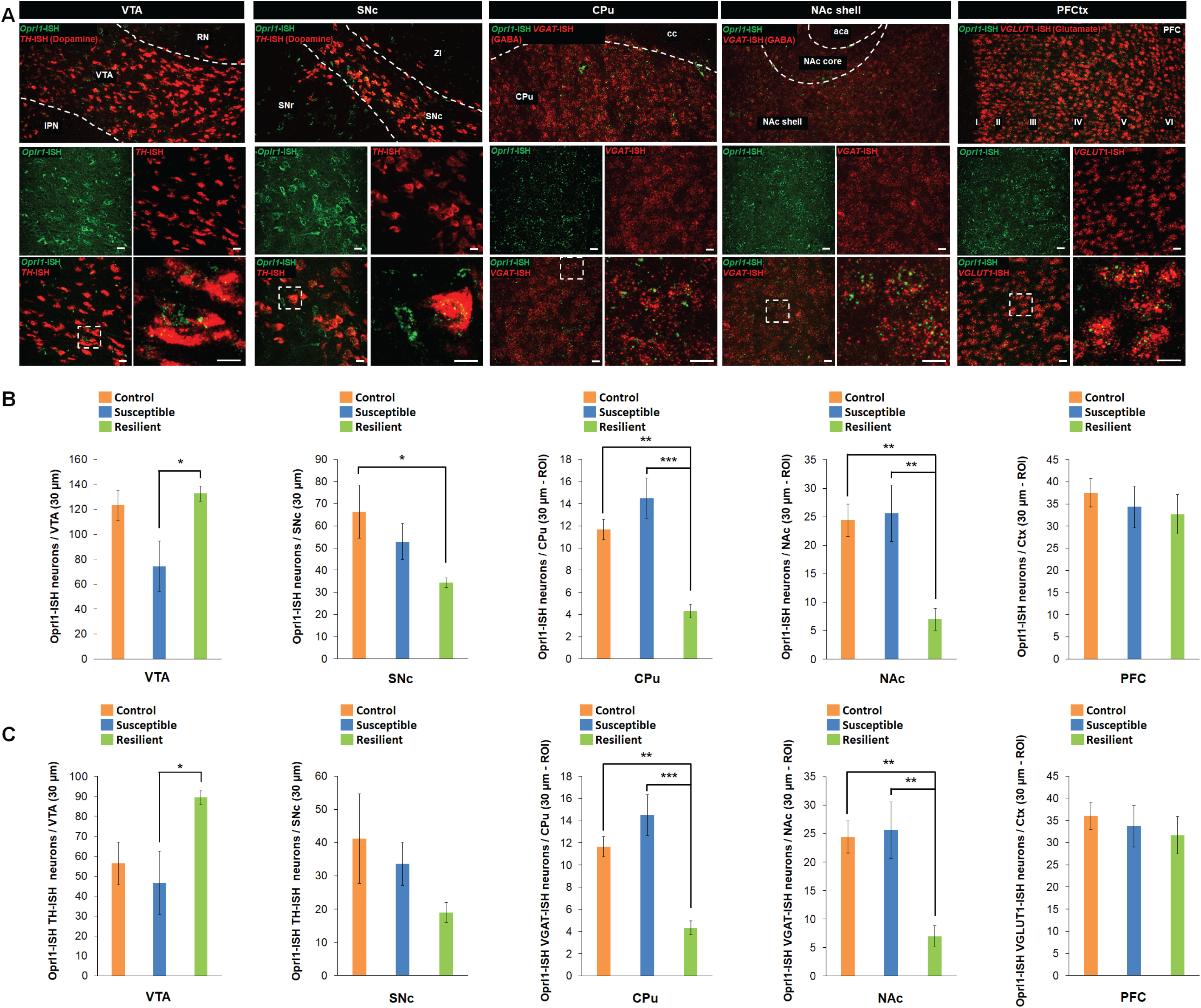
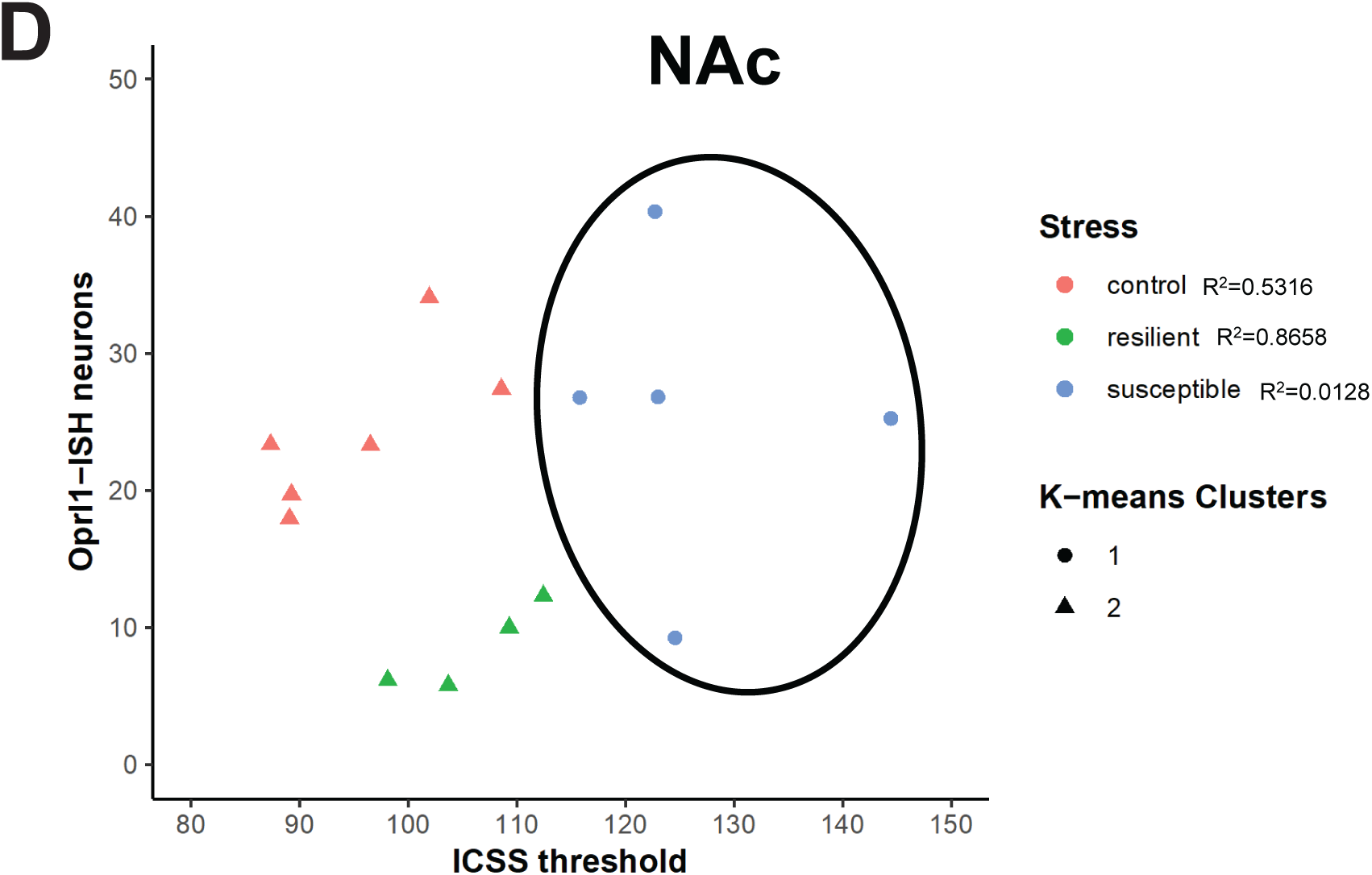
**(A)** Confocal images of *Oprl1* (green) RNAscope in-situ hybridization (*Oprl1*-ISH) in Ventral Tegmental Area (VTA), Substantia Nigra pars compacta (SNc), Caudate-Putamen (CPu), Nucleus Accumbens shell (NAc) and Prefrontal Cortex (PFC). The main NT-expressing cell type for each brain region was counterstained in red: Tyrosine Hydroxylase (TH) for VTA and SNc; Vescicular GABA transporter (VGAT) for CPu and NAc; Vescicular glutamate transporter 1 (*VGLUT1*-ISH) for PFC. **(B)** Quantification of neurons expressing *Oprl1* transcript in Control (N=6), susceptible (N=6) and resilient (N=6) rats for each brain region. ICSS-based clustering into susceptible and resilient groups revealed an heterogeneous regulation of *Oprl1*. The decrease in *Oprl1* + neurons is found mostly in resilient rats for NAc, and in susceptible rats in VTA. Tukey’s multiple comparisons, * p<0.05, ** p < 0.01, *** p < 0.001. **(C)** Quantification of NT-expressing *Pnoc*+ neurons (TH for VTA, SNc; VGAT in CPu, NAc; VGLUT1 in PFC) in Control (N=6), susceptible (N=6) and resilient (N=6) rats for each brain region. Tukey’s multiple comparisons, * p<0.05, ** p < 0.01, *** p < 0.001. **(D)** A data-driven unbiased clustering algorithm (k-means) that uses geometric distance between data points to separate data sets in k clusters predicted the susceptible group in the NAc with high accuracy when given k=2 and ICSS threshold and number of *Oprl1*+ neurons as variables to calculate distances.

### Study 2 Interim Discussion

Rats exposed to 21 days of social defeat exhibited marked alterations in the number of *Pnoc*-expressing neurons in the VTA, dorsal striatum, and PFC and parallel decreases occurred in postsynaptic receptor (*Oprl1*) expression. The VTA pattern replicated and extended the findings from Study 1 highlighting increased *Pnoc* expression in the VTA of both male and female mice. These changes might contribute to a mechanism by which chronic stress affects neuronal function through plasticity in neurotransmitter expression [55,56]. The resilient rats – despite being behaviorally similar to controls – appeared to actively regulate nociceptin signaling by downregulating *Pnoc* in VTA non-dopaminergic neurons, and postsynaptic *Oprl1* receptors in nucleus accumbens GABAergic neurons. Given that both *Pnoc* ligand and its *Oprl1* receptor mRNA are downregulated in these two regions involved in reward processing, and that N/OFQ signaling is generally inhibitory, these changes could result in increased activation of nucleus accumbens GABAergic medium spiny neurons (decrease of inhibition) and in turn boost GABAergic inhibition of the VTA and other targets.

Nociceptin receptors within the nucleus accumbens are thought to play a crucial role in the body’s stress response. These receptors exert their influence primarily by modulating dopamine signaling, thereby affecting encoding of both rewarding and aversive stimuli [57]. This modulation of dopamine release within the nucleus accumbens could dampen motivation and reinforced behaviors [58]. Although Study 2 confirmed that exposure to a chronic stressor altered the number of *Pnoc*-expressing neurons in the VTA, dorsal striatum, and PFC and reduced postsynaptic receptor (*Oprl1*) expression and induced anhedonic behavior among susceptible rats, it is unclear whether NOPR antagonism might affect more complex forms of anhedonic behaviors, including the animals’ propensity to modulate behavior as a function of reward. Using a translational task probing reward learning [59] that was modeled after similar tasks widely used in humans with MDD and anhedonia [46,47], the goal of Study 3 was to test the hypothesis that single administration of a highly selective NOPR antagonist (BTRX-246040) would boost reward learning in (non-stressed) rats.

### Study 3: Single Administration of a NOPR Antagonist Potentiates Reward Learning in Rats

#### Methods

##### Subjects

Male Long Evans rats (n = 8) obtained from Charles River Laboratories (Wilmington, MA) were housed in a climate-controlled vivarium with a 12-h light/dark cycle (lights on at 7am) and maintained at approximately 80% of their free-feeding weight via post-session portions of rodent chow. The protocol was approved by the Institutional Animal Care and Use Committee at McLean Hospital and in accordance with guidelines from the Committee on Care and Use of Laboratory Animals of the Institute of Laboratory Animals Resources, Commission on Life Sciences [60].

##### Apparatus

Details and schematics of the rat touch-sensitive experimental chamber can be found in [61]. Briefly, a Plexiglas chamber (25×30×35 cm) was situated in a sound- and light-attenuating enclosure (40×60×45 cm). A 17” touch-sensitive screen (1739L, ELO TouchSystems, Menlo Park, CA) comprised the inside right-hand wall of the chamber. An infusion pump (PHM-100-5, Med Associates, St. Albans, VT) outside the enclosure was used to deliver sweetened condensed milk solution (Sysco Corporation, Houston, TX) into the shallow reservoir (diameter: 3 cm) of a custom-designed aluminum receptacle (4×5×1 cm) that was mounted 2 cm above the floor bars and centered on the left-hand inside wall. A speaker bar (NQ576AT, Hewlett-Packard, Palo Alto, CA) mounted above the touchscreen was used to emit audible feedback. All experimental events and data collection were programmed in E-Prime Professional 2.0 (Psychology Software Tools, Inc., Sharpsburg, PA).

##### Procedure

*Line length discrimination training*. Modified response-shaping techniques were used to train rats to engage with the touchscreen [62]. A 5×5 cm blue square on a black background served as a response box and was centered on the touchscreen with its lower edge 10 cm above the floor bars. This required the rat to rear on its hind legs to make a touchscreen response with its paw. Each response was reinforced with 0.1 ml of 30% sweetened condensed milk, paired with an 880-ms yellow screen flash and a 440 Hz tone, and followed by a 5-sec blackout period. Following reliable responding by the rats, the position of the response box was alternated 5 cm left and right of center across 100-trial training sessions. After responses with latencies <5 sec were reliably observed to each position, line-length discrimination training commenced. Discrete trials began with presentation of a white line, with its lower edge presented 1.5 cm above the left and right response boxes. The length of the line was either 600 px (31.5 cm: long line) or 200 px (10.5 cm: short line); the width of both lines was 120 px (6.5 cm). Long and short line-length trial types varied in a quasi-random manner across 100-trial sessions such that there were exactly 50 trials of each type, but a given trial type would not be presented more than 5 times in a row. Subjects were trained to respond to the left or right response box depending on the length of the white line (long line: respond left, short line: respond right, or vice versa). Response box designation was counter-balanced across subjects. A correct response was reinforced as described above and was followed by a 5-sec blackout period, whereas an incorrect response immediately resulted in a 10-sec blackout period. A correction procedure was implemented during initial discrimination training, in which each incorrect trial was repeated until a correct response was made[63], and was discontinued after <10 repeats of each trial type occurred per session. Discrimination training sessions continued without correction until accuracies for both line length trial types were ≥80% correct for 3 consecutive sessions, concordant with the performance criterion of 75-85% correct in previous PRT studies with human subjects (refs). Following line length discrimination training, weekly (Mon-Fri) protocols were arranged such that sessions were conducted on Mon and Tues using the line-length stimuli described above in which all correct responses were rewarded [1:1 (100%:100%)]. The line length to be associated with the rich and lean contingency for the remainder of the week was determined for each subject during these 2 training sessions by examining their accuracies and designating the line length with a higher mean accuracy as the stimulus to be rewarded on the lean schedule. This was designed to examine the effects of the NOPR ligand BTRX-246040.

###### PRT Drug Tests

Following the establishment of probabilistic contingencies, a weekly (Mon-Fri) acute drug testing protocol was arranged as follows. Two maintenance sessions in which correct responses on all trials were reinforced (Mon and Tue) were followed by two sessions in which 3:1 (60%:20%) rich/lean probabilistic contingencies were arranged (Wed and Thurs). On the 5th session of the week (Fri), vehicle or a selected dose of BTRX-246040 (3, 10, or 30 mg/kg) was tested by administering it orally 60 min prior to a 3:1 (60%:20%) probabilistic session. Doses of BTRX-246040 were tested in a mixed order across subjects, and all doses of BTRX-246040 were tested in all subjects.

##### Drugs

BTRX-246040 was obtained from BlackThorn Therapeutics (San Francisco, CA) and was dissolved in a vehicle of 20% captisol and titrated to a pH of ∼2 before the addition of the drug. Drug doses were administered per os (p.o.) via gavage in a volume of 5 ml/kg, 60 min prior to the experimental session. Drug doses, vehicle, and pretreatment time were predetermined by BlackThorn Therapeutics.

#### Results

As shown in the top panel of Figure 4, administration of 3 and 10 mg/kg BTRX-246040 produced log b values of 0.28 (±0.05) and 0.27 (±0.11), respectively, that were similar to those observed following treatment with vehicle [0.33 (±0.09)]. Administration of 30 mg/kg BTRX-246040, however, increased the group mean log b to 0.57 (±0.08). A one-way repeated-measures ANOVA with Greenhouse-Geisser correction of the log b metric did not demonstrate a main effect of *Treatment* (*F*(2.1,15) = 2.80, *p* = 0.09). However, since this study included exploratory dose range finding, we conducted subsequent paired t-test analyses between the vehicle and 30 mg/kg BTRX-246040 group, and this test demonstrated a significant difference (*t*(7) = 1.90, one-tailed *p* = 0.05; Cohen’s *d* = 0.67]. As shown in the middle panel of Figure 4, log d values remained unperturbed following administration of vehicle and all doses of BTRX-246040 and this was confirmed statistically following both repeated-measures ANOVA (*F*(2.3,16) = 0.77, *p* = 0.49) and paired t-test analysis [vehicle vs 30 mg/kg BTRX-246040: *t*(7) = 0.29, *p* > 0.35; *d* = 0.10). Finally, as shown in the bottom panel of Figure 4, control analyses indicated that median reaction times were not altered in any systematic fashion across treatment conditions. Two-way repeated measures ANOVA using Greenhouse-Geisser correction indicated no main effect of treatment (*F*(2.37,33.15) = 0.58, *p* > 0.55], no main effect of stimulus type (rich vs lean; *F*[1,14] = 0.68, *p* > 0.40], and no significant interaction [*F*(3,42) = 1.28, *p* > 0.30).

**Figure 4.**
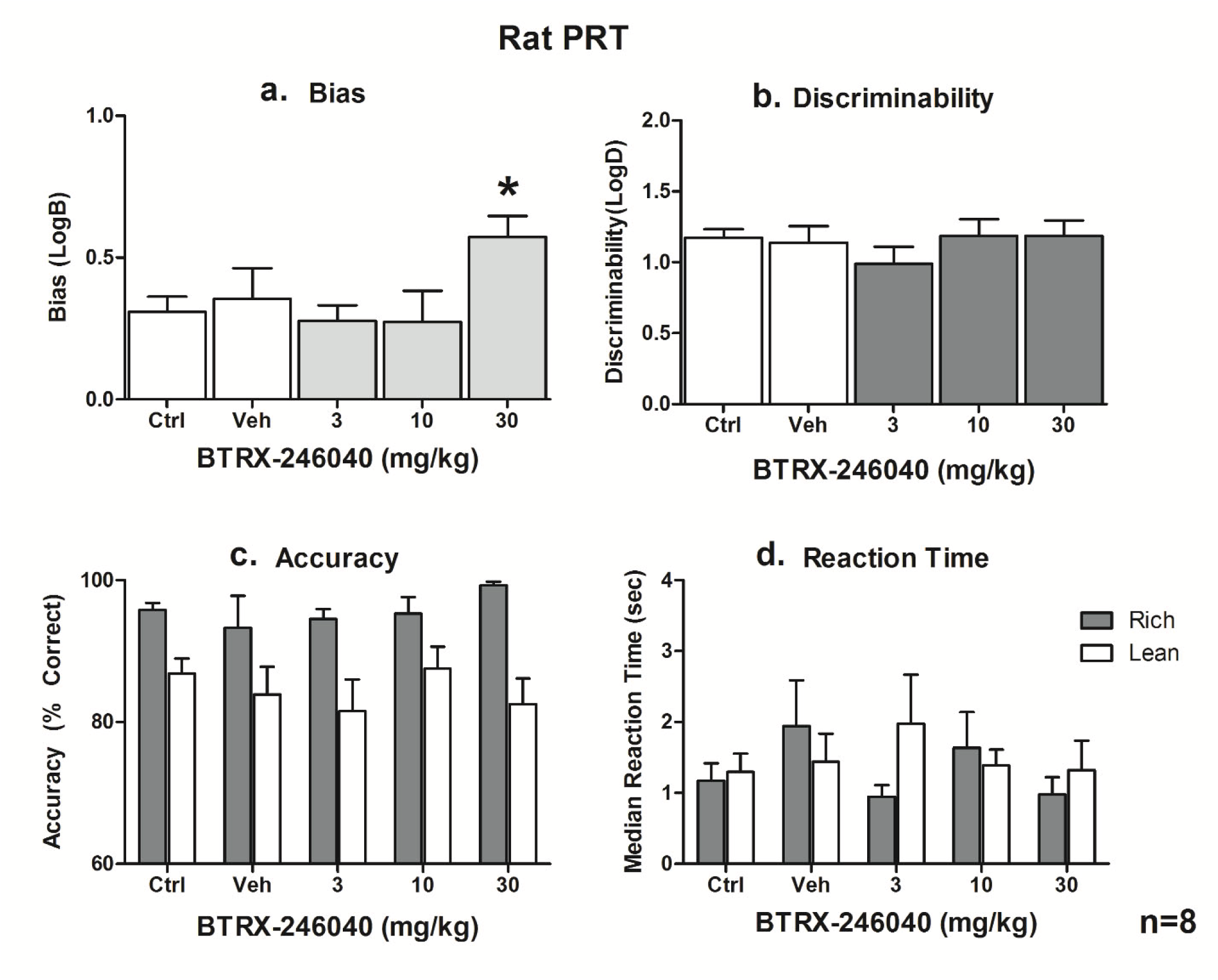
Effects of a single administration of BTRX-246040 in rats on **(A)** response bias (log b), (**B**) discriminability (log d), and **(C)** reaction time (group median).

### Study 3 Interim Discussion

These findings derived from a reverse-translated task that has been extensively used in humans with depression and anhedonia indicate that single administration of 30 mg/kg of a NOPR antagonist (BTRX-246040) improved reward responsiveness. Although a significant increase in log b values was only observed following the highest dose tested in this study, the magnitude of effect is large in the context of previous preclinical and clinical studies [46,47,64,65]. Moreover, as the doses tested were defined *a priori* (by Blackthorn Therapeutics), it is possible that higher doses might produce an even greater enhancement of reward sensitivity. Future studies of doses of BTRX-246040 higher than 30 mg/kg can help determine if this is the case. Having established that a single dose of a highly selective NOPR antagonist boosted reward learning in rats, in Study 4, we tested the novel hypothesis whether the human analogue of this compound increased motivation for rewards among individuals with MDD.

### Study 4: 8-week Administration of a NOPR Antagonist Potentiates Willingness to Exert Effort for Rewards in MDD

#### Methods

Data for these analyses were derived from a Phase 2a, 8-week, randomized, double-blind, placebo-controlled, parallel-group, multicenter study performed by Blackthorn Therapeutics (NEP-MDD-201; https://clinicaltrials.gov/study/NCT03193398) evaluating the efficacy and safety of BTRX-246040 (formerly known as LY2940094) – a selective nociceptin receptor antagonist – which had shown preclinical and clinical evidence of antidepressant/anti-anhedonic properties [19,66,67].

Patients were consented and screened for the study until approximately 100 patients were randomized. Randomization was performed at Visit 2 (baseline) in a 1:1 ratio to one of two treatment arms (BTRX-246040 or placebo) after meeting diagnostic criteria for MDD and study inclusion (see Supplement Material for a full list of the inclusion and exclusion criteria). Randomization was stratified by anhedonia symptom severity, as assessed by the Snaith Hamilton Pleasure Scale [68] (score ≤4 or >4). The BTRX-246040 starting dose for all eligible patients was 40 mg QD. A dose escalation to 80 mg QD occurred at Visit 3 (Week 1) visit for all patients. Patients who tolerated the dose increase remained at a dose of 80 mg QD for the subsequent 7 weeks. The dose could have been decreased back to 40 mg QD anytime through Visit 4 (Week 2), if required, for tolerability. Patients who could not tolerate 40 mg daily were discontinued from the study. Subsequent visits included Visit 5 (Week 4), Visit 6 (Week 6), and the end of the treatment period, Visit 7 (Week 8). All patients returned to the study site for a safety follow-up visit approximately 1 to 2 weeks after discontinuation of study drug. Overall duration in the study for each patient was approximately 14 weeks.

The primary endpoint – changes in clinician-administered Montgomery-Asberg Depression Rating Scale (MADRS) total score – was not met [69]. In light of preclinical evidence indicating that NOPR antagonism has pro-hedonic and motivational effects [67], in the current analyses, we evaluated whether, relative to placebo, 8-week treatment with BTRX-246040 boosted willingness to exert effort for rewards using the Effort Expenditure for Reward Task (EEfRT) [70].

##### Participants

Analyses were performed on individuals with MDD data who performed the EEfRT task and completed the post-treatment clinical assessments (N = 52 randomized to BTRX-246040 and 50 randomized to placebo) (Table 1, see also CONSORT diagram, Supplemental Figure 1).

**Table 1:**
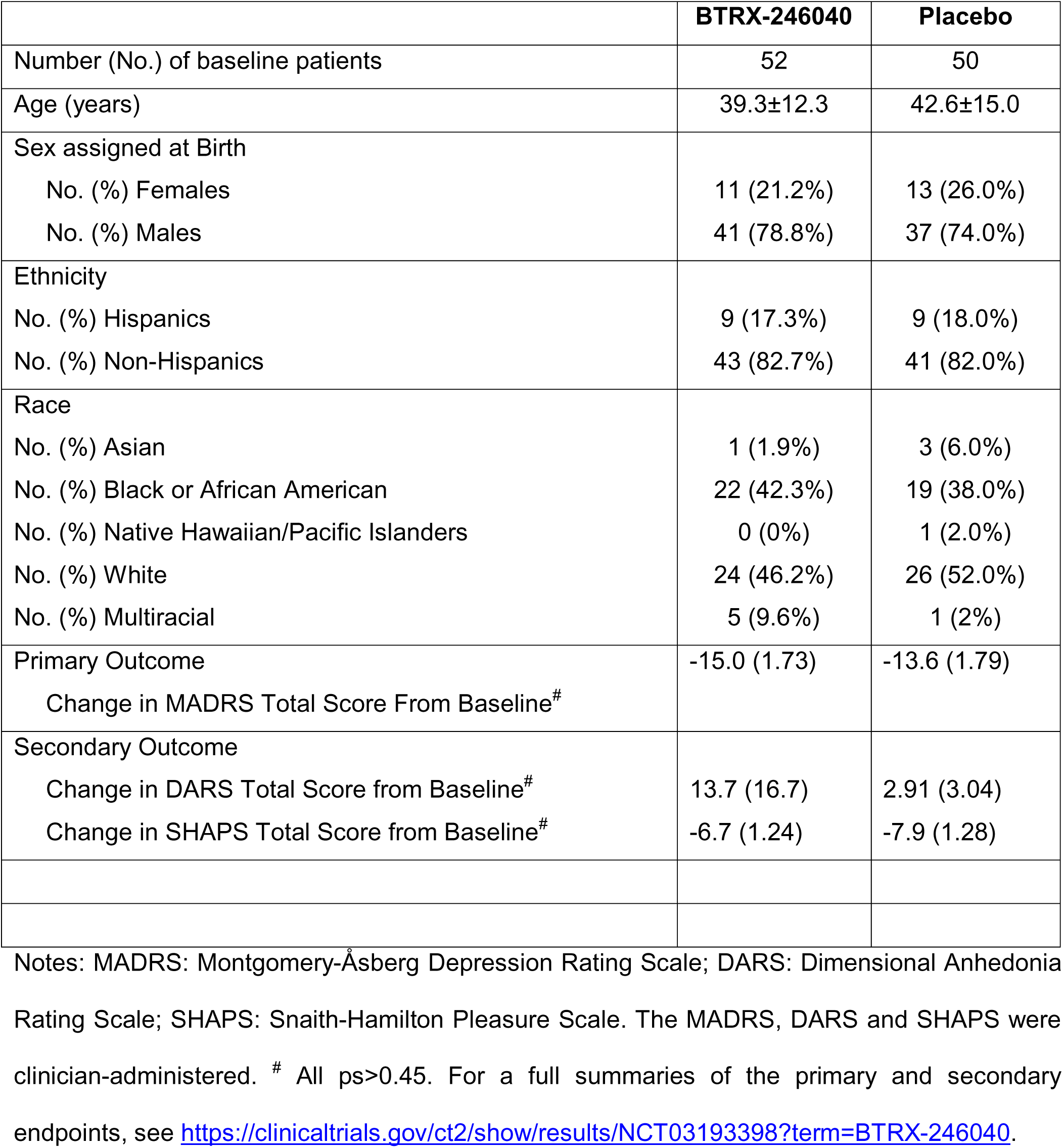
Sociodemographic and selected clinical variables.

|  | **BTRX-246040** | **Placebo** |
| --- | --- | --- |
| Number (No.) of baseline patients | 52 | 50 |
| Age (years) | 39.3±12.3 | 42.6±15.0 |
| Sex assigned at Birth |  |  |
| No. (%) Females | 11 (21.2%) | 13 (26.0%) |
| No. (%) Males | 41 (78.8%) | 37 (74.0%) |
| Ethnicity |  |  |
| No. (%) Hispanics | 9 (17.3%) | 9 (18.0%) |
| No. (%) Non-Hispanics | 43 (82.7%) | 41 (82.0%) |
| Race |  |  |
| No. (%) Asian | 1 (1.9%) | 3 (6.0%) |
| No. (%) Black or African American | 22 (42.3%) | 19 (38.0%) |
| No. (%) Native Hawaiian/Pacific Islanders | 0 (0%) | 1 (2.0%) |
| No. (%) White | 24 (46.2%) | 26 (52.0%) |
| No. (%) Multiracial | 5 (9.6%) | 1 (2%) |
| Primary Outcome | -15.0 (1.73) | -13.6 (1.79) |
| Change in MADRS Total Score From Baseline<sup>#</sup> |  |  |
| Secondary Outcome |  |  |
| Change in DARS Total Score from Baseline<sup>#</sup> | 13.7 (16.7) | 2.91 (3.04) |
| Change in SHAPS Total Score from Baseline<sup>#</sup> | -6.7 (1.24) | -7.9 (1.28) |

Notes: MADRS: Montgomery-Åsberg Depression Rating Scale; DARS: Dimensional Anhedonia Rating Scale; SHAPS: Snaith-Hamilton Pleasure Scale. The MADRS, DARS and SHAPS were clinician-administered. <sup>#</sup> All ps>0.45. For a full summaries of the primary and secondary endpoints, see https://clinicaltrials.gov/ct2/show/results/NCT03193398?term=BTRX-246040.

##### Clinical trial design

Patients randomized to BTRX-246040 received 40 mg administered orally for 1 week (1 capsule), followed by 80 mg (2 capsules) for 7 weeks, for a total of 8 weeks of treatment. Similarly, patients randomized to placebo received 1 capsule for 1 week, followed by 2 capsules for 7 weeks. Drug and placebo capsules were matched. Fifty-two patients (11 females; 39.3 ± 12.3 years old) were randomized to BTRX-246040, whereas 50 received placebo (13 females; 42.6 ± 15.0).

##### Clinical Scales

The 10-item version of the MADRS [71] served as the primary endpoint. Scores between 0 and 6 indicate no depression, between 7 and 19 mild depression, between 20 and 34 moderate depression, and between 35 and 60 severe depression. Among others, the Dimensional Anhedonia Rating Scale (DARS, [72]) and Snaith-Hamilton Pleasure Scale (SHAPS; [68]) served as secondary endpoints. These scales were investigator-administered (for other clinical endpoints, see https://clinicaltrials.gov/ct2/show/results/NCT03193398?term=BTRX-246040).

##### Effort Expenditure for Reward Task (EEfRT)

The EEfRT was developed to assess willingness to exert effort relative to size and likelihood of reward [70]). In each trial, participants are given an opportunity to choose between two different task difficulty levels in order to obtain monetary rewards: an easy choice requiring 30 keypresses using the index finger of their dominant hand delivering $1.00 as reward vs. a difficult choice requiring approximately 100 keypresses using the little finger of the non-dominant hand in order to receive larger rewards (range: $1.24-$4.12). Patients are required to complete each trial within a constrained period of time in order to obtain a monetary reward. The task was administered to the patient and reviewed by site personnel qualified to oversee completeness.

##### EEfRT Data Cleaning and Analysis

Data were initially evaluated for quality and cleaned to ensure that individuals did not time out more than 20% and did not fail to complete more than 20% of selected options for any EEfRT task administration. Of the initial 102 participants who were recruited, the data cleaning led to an exclusion of 6 participants (BTRX-246040: n=3; placebo: n=3). Consistent with prior studies, we examined both the total proportion of hard task choices selected at each session, as well as the model-derived parameters from computational models previously described [73–75]. For purposes of computational modeling for each subject we tested several variants of a subjective value (SV) model, as given by Equation 1:

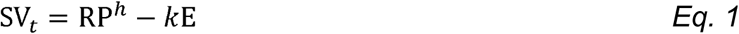

Where R is the reward available for the high effort option, P is the probability of winning, and E is the effort required. The free parameters h and k represent sensitivity to probability and effort information. Subjective value estimates were then transformed to choice probabilities using the Softmax function (Eq 3).

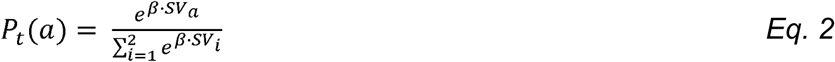

To account for the use of multiple repeated administrations, we fit several variants of this model that held two parameters constant across all three timepoints, and allowed one parameter to vary. This was done to achieve greater model fit (as indexed by BIC) as compared to a model where all three parameters were allowed to vary for each session. SV model 1 allowed the *k* parameter to vary, while holding *h* and *t* constant; SV model 2 allowed the *t* parameter to vary while holding *k* and *h* constant, and SV model 3 allowed the *h* parameter to vary while holding *k* and *t* constant. We then compared performance of these three models to a Bias model described by Equation 3. This model assumes that participants do not use reward value or reward probability information to guide effort expenditure. The probability of selecting the hard option under this model is computed as:

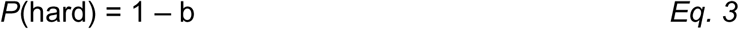

For purposes of model comparison to the SV models, we included a separate bias parameter for each time of the three timepoints.

##### Model fitting

To estimate model fits, we used maximum likelihood estimation as implemented via the fmincon function in Matlab. Options for fmincon were set at a max of 10,000 function evals and 4,000 iterations. For the k and h parameters of the SV model, upper and low bounds were set at 10 and 0, respectively. For the t parameter, upper and lower bounds were set at 100 and 0, respectively. For each model, a total of 100 random parameter initializations was used. Bayesian information criterion (BIC; [76]) was used to compare SV model fit to the Bias model.

##### Statistical Analysis

All analyses of group effects were tested using linear mixed effects models as implemented by the fitlme function in Matlab. For proportion of hard task choices, all participants with available data on two or more sessions were included. For analysis of model-derived effort discounting parameters, we only included individuals with complete data at all three timepoints, and individuals who were better fit by each of the three SV models (relative to the Bias model) for each of the three parameters (k, h and t) tested (see Table 2). For all models tested, age and sex were included as covariates.

**Table 2:**
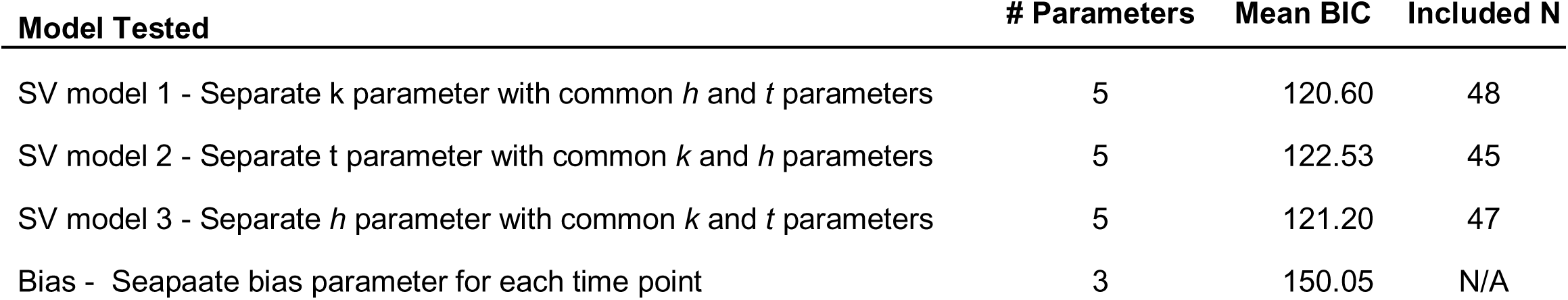
Summary of models that were evaluated.

#### Results

There was no significant interaction between *Treatment Arm* (BTRX-246040, placebo) and *Time* (baseline, week 4, week 8) for the total number of hard tasks selected. For the *k* discounting parameter, there was a trend-level interaction at week 7 (b = -.22, p = 0.063) such that the k discounting parameter was nominally increasing for the placebo group over time, but not for the BRTX-246060 group (see Figure 5A). For the inverse temperature (*t*) parameter, there was a significant interaction at week 7 (b = .84, p = 0.015) such that the inverse temperature was significantly increasing for the BRTX 246060 group (b = .88, p < .001), but not for the placebo group (b = .02, p = .944) (Figure 5B).

**Figure 5.**
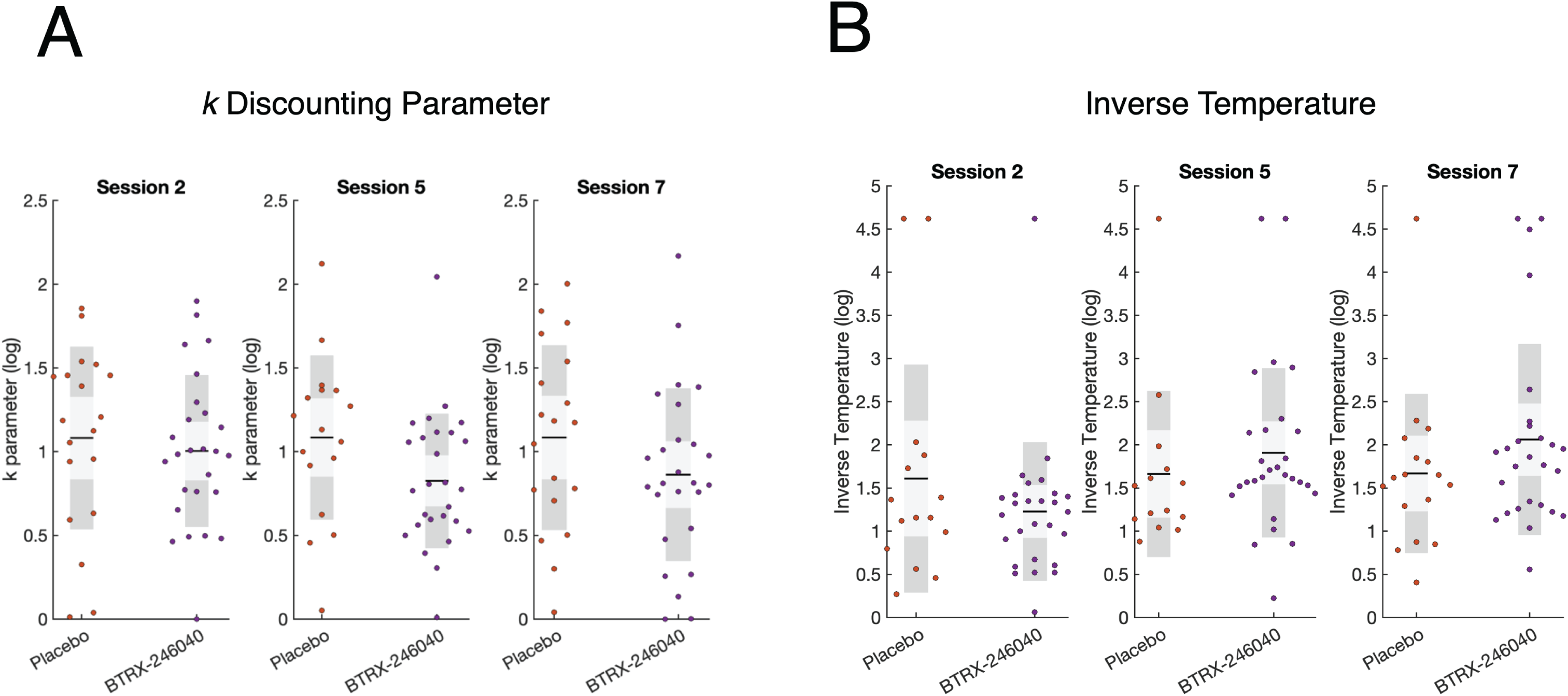
Effects of sustained BTRX-246040 treatment of EEfRT performance. **(A)** Relative to placebo, sustained treatment with BTRX-246040 was associated with a trend-level change in the *k* effort discounting parameter at week 8. **(B)** Relative to placebo, sustained treatment with BTRX-246040 was associated with a significant change in the *t* inverse temperature parameter at week 8.

### Study 4 Interim Discussion

The current results provide the first evidence in humans that 8-week treatment with a highly selective NOPR antagonist increased incentive motivation among individuals with MDD. Specifically, our results suggested that, relative to placebo, patients treated with an NOPR antagonist showed a margin trend towards being willing to expend consistent levels of effort over the course of three timepoints, and showed a marked increase in the consistency of their choices over time. Choice consistency as indexed by the inverse temperature parameters has frequently been observed as being abnormally lowered in MDD patients [77], and may reflect a lack of confidence in option valuation. Importantly, these effects were observed after excluding participants with poor data QA and poor model fits, suggesting they are unlikely to represent patterns of random responding or other sources of noise. It is also worth noting that these results emerged in the context of null findings with respect to the primary (MADRS scores) and various secondary (e.g., SHAPS scores, anxiety scale scores) endpoints (see https://clinicaltrials.gov/ct2/show/results/NCT03193398?term=BTRX-246040). Reasons for these dissociations are unclear but we speculate that the objective nature of the EEfRT, the focus on a precise and neurobiological well-characterized phenotype (incentive motivation), and the heterogenous nature of an “MDD” diagnosis might have contributed. Future studies, including those using the same NOPR antagonist in conjunction with neuroimaging, are warranted to evaluate whether NOPR antagonism might rescue reward-related neural abnormalities.

### General Discussion

Our understanding of the pathophysiology of MDD remains incomplete, which has contributed to modest innovation with respect to antidepressant treatments harnessing novel mechanisms. Here, across four studies that spanned three species (rats, mice and humans), evaluated different types of chronic stressor (early-life adversity, chronic social defeat), probed different brain regions implicated in the pathophysiology of MDD (striatum, PFC, VTA, cingulate), and used various endpoints to probe anhedonic phenotypes (sucrose preference, ICSS, reward learning, incentive motivation), we focused on the potential role of NOPR and its receptor in the emergence of stress-related depressive and anhedonic phenotypes. In addition, we tested the novel hypothesis that NOPR antagonism would boost two critical domains of the Research Domain Criteria (RDoC) Positive Valence Systems: reward learning and willingness to exert effort for rewards (incentive motivation).

Several notable findings emerged. First, exposure to both an early and later chronic stressor (LBN in Study 1 and chronic social defeat in Study 2) induced various anhedonic phenotypes (reduced sucrose preference in Study 1, altered ICSS reward threshold in Study 2). Consistent with our hypotheses, both types of stressors were associated with differential *Pnoc* expression in the VTA and striatum. Specifically, the early stressor (LBN) increased *Pnoc* expression in the VTA for both male and female mice, whereas it did so in the dorsal striatum for only female mice; thus, in the striatum, the effects showed sex- and region-specific selectivity. When considering adult rats exposed to chronic social defeat, we found that stressed rats were characterized by marked alterations in the number of *Pnoc*-expressing neurons in the VTA, dorsal striatum, and PFC and parallel decreases were observed in postsynaptic receptor (*Oprl1*) expression. Thus, the finding of increased *Pnoc* expression in the VTA was replicated in Study 1 and Study 2. Notably, resilient rats (rats not developing an anhedonic phenotype when facing the chronic social defeat) appeared to actively regulate N/OFQ signaling by downregulating *Pnoc* in VTA non-dopaminergic neurons, and postsynaptic *Oprl1* receptors in nucleus accumbens GABAergic neurons. Owing to the fact that (1) both *Pnoc* ligand and its *Oprl1* receptors are downregulated in these two regions that are important for reward processing and (2) *Pnoc* signaling is generally inhibitory, we speculate that the observed patterns might result in increased activation of nucleus accumbens GABAergic medium spiny neurons (decrease of inhibition) and in turn boost GABAergic inhibition of the VTA and other targets. Ultimately, this cascade of effects might contribute to the emergence of anhedonic phenotypes.

Finally, consistent with prior preclinical evidence highlighting that NOPR antagonism has anti-depressant and anti-anhedonic effects [3,19,78], we found that single administration of 30 mg/kg of BTRX-246040 – a highly selective and potent NOPR antagonist, which has 3- to 10-fold higher potency than the standard NOPR antagonist SB-612111 [15] – boosted rats’ ability to modulate behavior as a function of reward (they developed a stronger response bias toward a more frequently rewarded stimulus). Critically, we extended this finding in rats by showing that 8-week treatment with BTRX-246040 was associated with significantly higher willingness to exert efforts to pursue high rewards among humans with MDD. Of note, both reward learning and reward motivation have been found to be strongly dependent on dopaminergic signaling [34,35]. These findings are consistent with substantial *in vitro* and *in vivo* evidence pointing to an inhibitory effect of endogenous N/OFQergic signaling on monoaminergic (dopaminergic) neurotransmission in regions relevant to MDD. In line with this, NOPR has been found to inhibit tyrosine hydroxylase phosphorylation, and thus DA transmission presynaptically [79], and inhibit dopamine release in the VTA, nucleus accumbens and caudate nucleus [18–20]. Moreover, NOPR are found on the cell bodies of dopamine neurons in the midbrain and NOPR mRNA has been co-localized with tyrosine neurons as well as tegmental and nigral DA neurons [27]. Conversely, NOPR antagonism was found to enhance DA transmission [32,33], in addition to exerting antidepressant and anti-anhedonic effects [9–13,18,19]. Altogether, these data suggest that NOPR antagonists might exert their antidepressant effects by counteracting the inhibitory effects of the endogenous NOPR on monoaminergic pathways both pre- and post-synaptically. The current cross-species findings emerging from Study 3 and 4 are intriguing in light of preclinical evidence that BTRX-246040 had rapid-onset effects in various depression-relevant assays [12,17,18]. Thus, future studies in humans with MDD and/or anhedonia are warranted to evaluate whether BTRX-246040 could rapidly normalize anhedonic phenotypes.

Although the current work has several strengths, including integration across several species, approaches, and functional domains, a few limitations should be emphasized. First, in some studies, we did not assess female animals or did not have sufficient power to formally evaluate possible sex-specific effects. In light of sex differences in the prevalence of MDD, future studies powered to assess for sex effects are needed. Second, in the human Phase 2 study, evidence of improvements in incentive motivation emerged in the context of null findings when considering multiple clinical scales assessing depression severity, anhedonia, and anxiety. Thus, it is unclear whether the current EEfRT findings point to increased sensitivity when assessing a neurobiologically well-characterized construct with substantial biological validity or rather possible false positive findings. Although we believe that the latter option is improbable in light of the extensive preclinical literature suggesting pro-hedonic effects with NOPR antagonism, replications are clearly needed. Despite these limitations, we present compelling evidence of an important role of the N/OFQ-NOPR system in the pathophysiology of depressive phenotypes. Given their significant impact on reward and stress responses, NOPR antagonists should be further evaluated as promising treatment for anhedonia and stress-related disorders, including MDD.

## Acknowledgments

These studies were supported by National Institute of Mental Health P50 MH119467 (Study 1-3; awarded to DAP), R37 MH068376 (Study 1 and 3); awarded to DAP), and National Institute of Neurological Disorders and Stroke R01 NS057690 (Study 2; awarded to DD). BDK was supported in part by National Institute on Drug Abuse R01-DA047575. The authors are solely responsible for the design and conduct of these studies, including analysis, and interpretation of the data; preparation, review, and approval of the manuscript; and decision to submit the manuscript for publication. The authors are grateful to Jane Tiller, FRCPsych, MBA, MPhil (former Chief Medical Officer, BlackThorn Therapeutics) for her critical contributions to Study 4 and to Tanya Wallace, PhD (former Senior Director, Head of Biology, BlackThorn Therapeutics) for her contribution to Study 3.

## Supplemental Material

### Study 4 Inclusion Criteria

- Patients (age range: 18-65 years old) with a diagnosis of MDD as defined by DSM-5 criteria and have had at least 1 prior major depressive episode in the past 10 years
- Patients must present with a new current episode of MDD and the duration of the current episode must be at least 4 weeks but not longer than 18 months.
- At Visit 1 (screening) and Visit 2 (baseline), patients must have clinically significant depressive symptoms defined by tandem (investigator- and computer-administered) Montgomery-Asberg Depression Rating Scale (MADRS) total scores ≥ 26 with a difference of ≤ 7 points between the Investigator- and computer-administered MADRS total scores
- Patients must have a CGI-S score ≥ 4 at Visit 2 (baseline).

### Study 4 Exclusion Criteria

- Patients who present with any current DSM-5 disorder other than MDD which is the focus of treatment.
- Patients who are homicidal in the opinion of the Investigator or are at suicidal risk (any suicide attempts within 12 months prior to Visit 1 [screening] or any suicidal intent, including a plan, within 3 months prior to Visit 1 [screening]; C-SSRS answer of “YES” on item 4 or 5 [suicidal ideation]; Investigator- or computer-administered MADRS score of ≥ 5 on item 10 [suicidal thoughts]; by Investigator clinical evaluation).
- Patients cannot have any history of substance or alcohol use disorder within 12 months prior to Visit 1 (screening) per DSM-5 criteria
- Patients must not have a clinically significant comorbid disease.

**Supplemental Figure 1:**
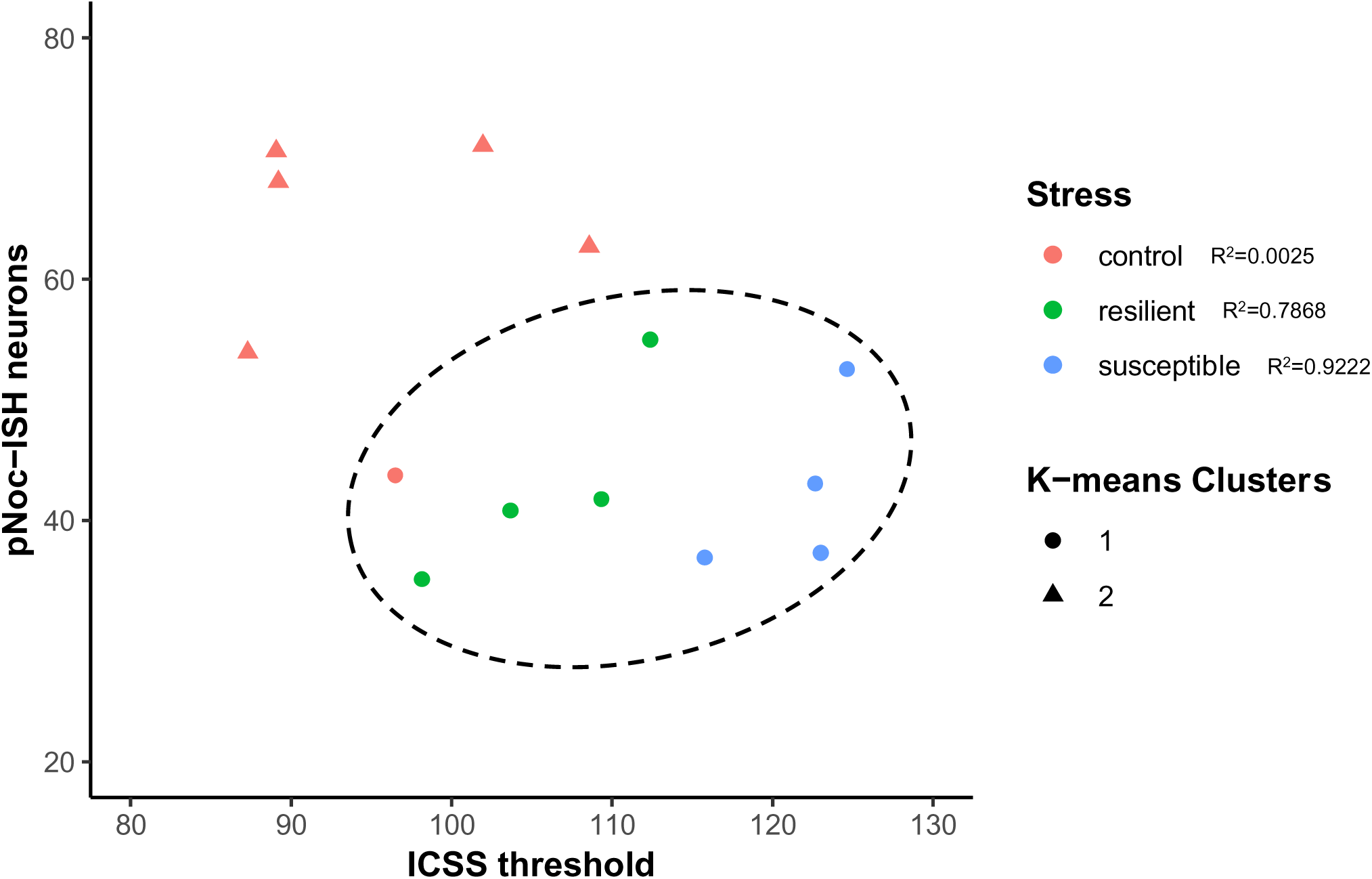
A data-driven unbiased clustering algorithm (k-means) that uses geometric distance between data points to separate data sets in k clusters predicted the split between control and stressed animals in the VTA with high accuracy when given k=2 and ICSS threshold and number of *Pnoc*+ neurons as variables to calculate distances.

**Supplemental Figure 2:**
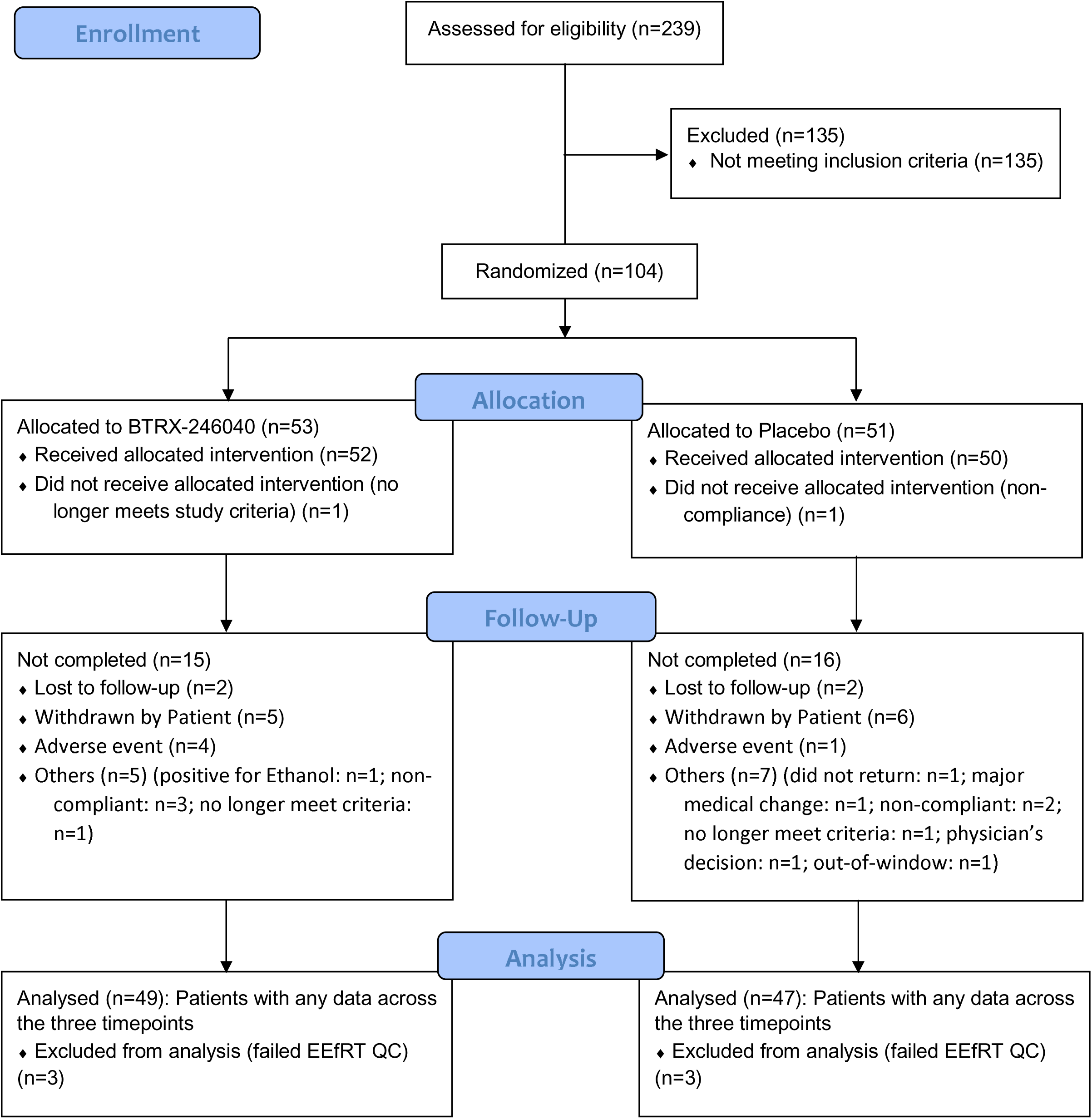
Consort diagram of patient flow.

## References

1. Gavioli EC, Holanda VAD, Ruzza C. NOP ligands for the treatment of anxiety and mood disorders. Handb Exp Pharmacol, 2018. p. 2018 Dec 8. doi: 10.1007/164_2018_188. [Epub ahead.

2. Witkin JM, Wallace TL, Martin WJ. Therapeutic approaches for NOP receptor antagonists in neurobehavioral disorders: Clinical studies in Major Depressive Disorder and Alcohol Use Disorder with BTRX-246040 (LY2940094). Handb Exp Pharmacol, 2018.

3. Ubaldi M, Cannella N, Borruto AM, Petrella M, Micioni Di Bonaventura MV, Soverchia L, et al. Role of Nociceptin/Orphanin FQ-NOP receptor system in the regulation of stress-related disorders. Int J Mol Sci. 2021;22.

4. Agorastos A, Chrousos GP. The neuroendocrinology of stress: the stress-related continuum of chronic disease development. Mol Psychiatry. 2022;27:502–513.

5. Pizzagalli DA. Depression, stress, and anhedonia: toward a synthesis and integrated model. Annu Rev Clin Psychol. 2014;10:393–423.

6. IsHak WW, Wen RY, Naghdechi L, Vanle B, Dang J, Knosp M, et al. Pain and Depression: A Systematic Review. Harv Rev Psychiatry. 2018;26:352–363.

7. Nativio P, Pascale E, Maffei A, Scaccianoce S, Passarelli F. Effect of stress on hippocampal nociceptin expression in the rat. Stress. 2012;15:378–384.

8. Der-Avakian A, D’Souza MS, Potter DN, Chartoff EH, Carlezon WA, Pizzagalli DA, et al. Social defeat disrupts reward learning and potentiates striatal nociceptin/orphanin FQ mRNA in rats. Psychopharmacology (Berl). 2017;234:1603–1614.

9. Gavioli EC, Marzola G, Guerrini R, Bertorelli R, Zucchini S, De Lima TCM, et al. Blockade of nociceptin/orphanin FQ-NOP receptor signalling produces antidepressant-like effects: pharmacological and genetic evidences from the mouse forced swimming test. European Journal of Neuroscience. 2003;17:1987–1990.

10. Rizzi A, Gavioli EC, Marzola G, Spagnolo B, Zucchini S, Ciccocioppo R, et al. Pharmacological characterization of the nociceptin/orphanin FQ receptor antagonist SB-612111 [(-)-cis-1-methyl-7-[[4-(2,6-dichlorophenyl)piperidin-1-yl]methyl]-6,7,8,9-tetrahydro-5H-benzocyclohepten-5-ol]: in vivo studies. Journal of Pharmacology and Experimental Therapeutics. 2007;321:968–974.

11. Redrobe J, Calo’ G, Regoli D, Quirion R. Nociceptin receptor antagonists display antidepressant-like properties in the mouse forced swimming test. Naunyn Schmiedebergs Arch Pharmacol. 2002;365:164–167.

12. Holanda VAD, Medeiros IU, Asth L, Guerrini R, Calo’ G, Gavioli EC. Antidepressant activity of nociceptin/orphanin FQ receptor antagonists in the mouse learned helplessness. Psychopharmacology (Berl). 2016;233:2525–2532.

13. Holanda VAD, Oliveira MC, Da Silva Junior ED, Calo’ G, Ruzza C, Gavioli EC. Blockade of nociceptin/orphanin FQ signaling facilitates an active copying strategy due to acute and repeated stressful stimuli in mice. Neurobiol Stress. 2020;13.

14. Vitale G, Ruggieri V, Filaferro M, Frigeri C, Alboni S, Tascedda F, et al. Chronic treatment with the selective NOP receptor antagonist [Nphe1,Arg14,Lys15]N/OFQ-NH2 (UFP-101) reverses the behavioural and biochemical effects of unpredictable chronic mild stress in rats. Psychopharmacology (Berl). 2009;207:173–189.

15. Ferrari F, Rizzo S, Ruzza C, Calo G. Detailed In Vitro Pharmacological Characterization of the Clinically Viable Nociceptin/Orphanin FQ Peptide Receptor Antagonist BTRX-246040. J Pharmacol Exp Ther. 2020;373:34–43.

16. Toledo MA, Pedregal C, Lafuente C, Diaz N, Martinez-Grau MA, Jiménez A, et al. Discovery of a novel series of orally active nociceptin/orphanin FQ (NOP) receptor antagonists based on a dihydrospiro(piperidine-4,7’-thieno[2,3-c]pyran) scaffold. J Med Chem. 2014;57:3418–3429.

17. D’Oliveira da Silva F, Azevedo Neto J, Sturaro C, Guarino A, Robert C, Gavioli EC, et al. The NOP antagonist BTRX-246040 increases stress resilience in mice without affecting adult neurogenesis in the hippocampus. Neuropharmacology. 2022;212.

18. Li Z, Xu Y, Li R, Sheng Z, Chen X, Liu X, et al. BTRX-246040 Acts Through the Ventrolateral Periaqueductal Gray to Exert Antidepressant-Relevant Actions in Mice. Int J Neuropsychopharmacol. 2023;26:483–495.

19. Post A, Smart TS, Krikke-Workel J, Dawson GR, Harmer CJ, Browning M, et al. A Selective Nociceptin Receptor Antagonist to Treat Depression: Evidence from Preclinical and Clinical Studies. Neuropsychopharmacology. 2016;41:1803–1812.

20. Witkin JM, Rorick-Kehn LM, Benvenga MJ, Adams BL, Gleason SD, Knitowski KM, et al. Preclinical findings predicting efficacy and side-effect profile of LY2940094, an antagonist of nociceptin receptors. Pharmacol Res Perspect. 2016;4:e00275.

21. Post A, Smart TS, Krikke-Workel J, Dawson GR, Harmer CJ, Browning M, et al. A Selective Nociceptin Receptor Antagonist to Treat Depression: Evidence from Preclinical and Clinical Studies. Neuropsychopharmacology. 2016;41:1803–1812.

22. Neal CR, Mansour A, Reinscheid R, Nothacker H-P, Civelli O, Akil H, et al. Opioid receptor-like (ORL1) receptor distribution in the rat central nervous system: Comparison of ORL1 receptor mRNA expression with125I-[14Tyr]-orphanin FQ binding. J Comp Neurol. 1999;412:563–605.

23. Sim LJ, Childers SR. Anatomical distribution of mu, delta, and kappa opioid- and nociceptin/orphanin FQ-stimulated [35S]Guanylyl-5?-O-(?-Thio)-triphosphate binding in guinea pig brain. J Comp Neurol. 1997;386:562–572.

24. Berthele A, Platzer S, Dworzak D, Schadrack J, Mahal B, Büttner A, et al. [3h]-nociceptin ligand-binding and nociceptin opioid receptor mrna expression in the human brain. Neuroscience. 2003;121:629–640.

25. Lohith TG, Zoghbi SS, Morse CL, Araneta MDF, Barth VN, Goebl NA, et al. Retest imaging of [11C]NOP-1A binding to nociceptin/orphanin FQ peptide (NOP) receptors in the brain of healthy humans. Neuroimage. 2014;87:89–95.

26. Mollereau C, Mouledous L. Tissue distribution of the opioid receptor-like (ORL1) receptor. Peptides (NY). 2000;21:907–917.

27. Norton CS, Neal CR, Kumar S, Akil H, Watson SJ. Nociceptin/orphanin FQ and opioid receptor-like receptor mRNA expression in dopamine systems. J Comp Neurol. 2002;444:358–368.

28. Murphy NP, Maidment NT. Orphanin FQ/nociceptin modulation of mesolimbic dopamine transmission determined by microdialysis. J Neurochem. 1999;73:179–186.

29. Murphy NP, Ly HT, Maidment NT. Intracerebroventricular orphanin FQ/nociceptin suppresses dopamine release in the nucleus accumbens of anaesthetized rats. Neuroscience. 1996;75:1–4.

30. Koizumi M, Midorikawa N, Takeshima H, Murphy NP. Exogenous, but not endogenous nociceptin modulates mesolimbic dopamine release in mice. J Neurochem. 2004;89:257– 263.

31. Arcuri L, Viaro R, Bido S, Longo F, Calcagno M, Fernagut PO, et al. Genetic and pharmacological evidence that endogenous nociceptin/orphanin FQ contributes to dopamine cell loss in Parkinson’s disease. Neurobiol Dis. 2016;89:55–64.

32. Marti M, Mela F, Fantin M, Zucchini S, Brown JM, Witta J, et al. Blockade of nociceptin/orphanin FQ transmission attenuates symptoms and neurodegeneration associated with Parkinson’s disease. J Neurosci. 2005;25:9591–9601.

33. Marti M, Mela F, Veronesi C, Guerrini R, Salvadori S, Federici M, et al. Blockade of nociceptin/orphanin FQ receptor signaling in rat substantia nigra pars reticulata stimulates nigrostriatal dopaminergic transmission and motor behavior. J Neurosci. 2004;24:6659–6666.

34. Kangas BD, Der-Avakian A, Pizzagalli DA. Probabilistic Reinforcement Learning and Anhedonia. Curr Top Behav Neurosci. 2022. 2022. 10.1007/7854_2022_349.

35. Treadway MT, Salamone JD. Vigor, Effort-Related Aspects of Motivation and Anhedonia. Curr Top Behav Neurosci. 2022. 2022. 10.1007/7854_2022_355.

36. Gavioli EC, Calo’ G. Nociceptin/orphanin FQ receptor antagonists as innovative antidepressant drugs. Pharmacol Ther. 2013;140:10–25.

37. Parker KE, Gomez AM, Pedersen CE, Spangler SM, Walicki MC, Feng SY, et al. A paranigral VTA nociceptin circuit that constrains motivation for reward. Cell.

38. Iturra-Mena AM, Kangas BD, Pizzagalli DA. Nociceptin Receptor Antagonism Modulates Electrophysiological Markers of Reward Learning. Int J Neuropsychopharmacol. 2023;26:496–500.

39. Ho TC, King LS. Mechanisms of neuroplasticity linking early adversity to depression: developmental considerations. Transl Psychiatry. 2021;11.

40. Pechtel P, Pizzagalli DA. Effects of early life stress on cognitive and affective function: an integrated review of human literature. Psychopharmacology (Berl). 2011;214:55–70.

41. Hanson JL, Williams A V., Bangasser DA, Peña CJ. Impact of Early Life Stress on Reward Circuit Function and Regulation. Front Psychiatry. 2021;12.

42. Pechtel P, Pizzagalli DA. Disrupted reinforcement learning and maladaptive behavior in women with a history of childhood sexual abuse: a high-density event-related potential study. JAMA Psychiatry. 2013;70:499–507.

43. Hueske E, Stine C, Yoshida T, Crittenden JR, Gupta A, Johnson JC, et al. Developmental and adult striatal patterning of nociceptin ligand marks striosomal population with direct dopamine projections. Journal of Comparative Neurology. 2024;532.

44. Lazaridis I, Crittenden JR, Ahn G, Hirokane K, Wickersham IR, Yoshida T, et al. Striosomes control dopamine via dual pathways paralleling canonical basal ganglia circuits. Current Biology. 2024;34:5263–5283.e8.

45. Evans RC, Twedell EL, Zhu M, Ascencio J, Zhang R, Khaliq ZM. Functional dissection of basal ganglia inhibitory iInputs onto substantia nigra dopaminergic neurons. Cell Rep. 2020;32:108156.

46. Pizzagalli DA, Jahn AL, O’Shea JP. Toward an objective characterization of an anhedonic phenotype: a signal-detection approach. Biol Psychiatry. 2005;57:319–327.

47. Pizzagalli DA, Iosifescu D, Hallett LA, Ratner KG, Fava M. Reduced hedonic capacity in major depressive disorder: evidence from a probabilistic reward task. J Psychiatr Res. 2008;43:76–87.

48. Walker CD, Bath KG, Joels M, Korosi A, Larauche M, Lucassen PJ, et al. Chronic early life stress induced by limited bedding and nesting (LBN) material in rodents: critical considerations of methodology, outcomes and translational potential. Stress. 2017;20:421–448.

49. Wendel KM, Short AK, Noarbe BP, Haddad E, Palma AM, Yassa MA, et al. Early life adversity in male mice sculpts reward circuits. Neurobiol Stress. 2021;15.

50. Molet J, Heins K, Zhuo X, Mei YT, Regev L, Baram TZ, et al. Fragmentation and high entropy of neonatal experience predict adolescent emotional outcome. Transl Psychiatry. 2016;6.

51. Bolton JL, Ruiz CM, Rismanchi N, Sanchez GA, Castillo E, Huang J, et al. Early-life adversity facilitates acquisition of cocaine self-administration and induces persistent anhedonia. Neurobiol Stress. 2018;8:57–67.

52. Bolton JL, Molet J, Regev L, Chen Y, Rismanchi N, Haddad E, et al. Anhedonia Following Early-Life Adversity Involves Aberrant Interaction of Reward and Anxiety Circuits and Is Reversed by Partial Silencing of Amygdala Corticotropin-Releasing Hormone Gene. Biol Psychiatry. 2018;83:137–147.

53. Kangas BD, Short AK, Luc OT, Stern HS, Baram TZ, Pizzagalli DA. A cross-species assay demonstrates that reward responsiveness is enduringly impacted by adverse, unpredictable early-life experiences. Neuropsychopharmacology. 2022;47:767–775.

54. Kangas BD, Ang YS, Short AK, Baram TZ, Pizzagalli DA. Computational Modeling Differentiates Learning Rate From Reward Sensitivity Deficits Produced by Early-Life Adversity in a Rodent Touchscreen Probabilistic Reward Task. Biological Psychiatry Global Open Science. 2024;4.

55. Prakash N, Stark CJ, Keisler MN, Luo L, Der-Avakian A, Dulcis D. Serotonergic Plasticity in the Dorsal Raphe Nucleus Characterizes Susceptibility and Resilience to Anhedonia. J Neurosci. 2020;40:569–584.

56. Dulcis D, Spitzer NC. Reserve pool neuron transmitter respecification: Novel neuroplasticity. Dev Neurobiol. 2012;72:465–474.

57. Soares-Cunha C, de Vasconcelos NAP, Coimbra B, Domingues AV, Silva JM, Loureiro-Campos E, et al. Nucleus accumbens medium spiny neurons subtypes signal both reward and aversion. Mol Psychiatry. 2020;25:3241–3255.

58. Khan MS, Boileau I, Kolla N, Mizrahi R. A systematic review of the role of the nociceptin receptor system in stress, cognition, and reward: relevance to schizophrenia. Transl Psychiatry. 2018;8.

59. Kangas BD, Wooldridge LM, Luc OT, Bergman J, Pizzagalli DA. Empirical validation of a touchscreen probabilistic reward task in rats. Transl Psychiatry. 2020;10.

60. National Research Council. Guidelines for the Care and Use of Laboratory Animals. 2011.

61. Kangas BD, Bergman J. Touchscreen technology in the study of cognition-related behavior. Behavioural Pharmacology. 2017;28:623–629.

62. Kangas BD, Bergman J. A novel touch-sensitive apparatus for behavioral studies in unrestrained squirrel monkeys. J Neurosci Methods. 2012;209:331–336.

63. Kangas BD, Branch MN. Empirical validation of a procedure to correct position and stimulus biases in matching-to-sample. J Exp Anal Behav. 2008;90:103–112.

64. Der-Avakian A, D’Souza MSS, Pizzagalli DAA, Markou A. Assessment of reward responsiveness in the response bias probabilistic reward task in rats: implications for cross-species translational research. Transl Psychiatry. 2013;3:e297.

65. Der-Avakian A, D’Souza MS, Potter DN, Chartoff EH, Carlezon WA, Pizzagalli DA, et al. Social defeat disrupts reward learning and potentiates striatal nociceptin/orphanin FQ mRNA in rats. Psychopharmacology (Berl). 2017;234:1603–1614.

66. Dawson GR, Post A, Smart TS, Browning M, Harmer CJ. Accuracy in recognising happy facial expressions is associated with antidepressant response to a NOP receptor antagonist but not placebo treatment. J Psychopharmacol. 2021;35:1473–1478.

67. Witkin JM, Wallace TL, Martin WJ. Therapeutic approaches for NOP receptor antagonists in neurobehavioral disorders: Clinical studies in Major Depressive Disorder and Alcohol Use Disorder with BTRX-246040 (LY2940094). Handb Exp Pharmacol, 2018.

68. Snaith RP, Hamilton M, Morley S, Humayan A, Hargreaves D, Trigwell P. A scale for the assessment of hedonic tone the Snaith-Hamilton Pleasure Scale. Br J Psychiatry. 1995;167:99–103.

69. Jane Tiller, Clark Gao, Yuelu Liu, Humberto Gonzalez, Parvez Ahammad. A novel explainable AI approach to differentiate drug and placebo responders: Offering the opportunity for enhanced patient selection. International Society for CNS Clinical Trials and Methodology (ISCTM), Copenhagen, Denmark, 5-7 September, 2019 ; 2019.

70. Treadway MT, Buckholtz JW, Schwartzman AN, Lambert WE, Zald DH. Worth the ‘EEfRT’? The effort expenditure for rewards task as an objective measure of motivation and anhedonia. PLoS One. 2009;4:e6598.

71. Montgomery SA, Asberg M. A new depression scale designed to be sensitive to change. Br J Psychiatry. 1979;134:382–389.

72. Rizvi SJ, Quilty LC, Sproule BA, Cyriac A, Bagby RM, Kennedy SH. Development and validation of the Dimensional Anhedonia Rating Scale (DARS) in a community sample and individuals with major depression. Psychiatry Res. 2015;229:109–119.

73. Whitton AE, Cooper JA, Merchant JT, Treadway MT, Lewandowski KE. Using Computational Phenotyping to Identify Divergent Strategies for Effort Allocation Across the Psychosis Spectrum. Schizophr Bull. 2024;50:1127–1136.

74. Soder HE, Cooper JA, Lopez-Gamundi P, Hoots JK, Nunez C, Lawlor VM, et al. Dose-response effects of d-amphetamine on effort-based decision-making and reinforcement learning. Neuropsychopharmacology. 2021;46:1078–1085.

75. Cooper JA, Barch DM, Reddy LF, Horan WP, Green MF, Treadway MT. Effortful goal-directed behavior in schizophrenia: Computational subtypes and associations with cognition. J Abnorm Psychol. 2019;128:710–722.

76. G S. ‘Estimating the Dimension of a Model.’ The Annals of Statistics. 1978;6:461–464.

77. Pike AC, Robinson OJ. Reinforcement Learning in Patients With Mood and Anxiety Disorders vs Control Individuals: A Systematic Review and Meta-analysis. JAMA Psychiatry. 2022;79:313–322.

78. Witkin JM, Wallace TL, Martin WJ. Therapeutic approaches for NOP receptor antagonists in neurobehavioral disorders: Clinical studies in Major Depressive Disorder and alcohol use disorder with BTRX-246040 (LY2940094). Handb Exp Pharmacol. 2019. 31 January 2019. 10.1007/164_2018_186.

79. Olianas MC, Dedoni S, Boi M, Onali P. Activation of nociceptin/orphanin FQ-NOP receptor system inhibits tyrosine hydroxylase phosphorylation, dopamine synthesis, and dopamine D(1) receptor signaling in rat nucleus accumbens and dorsal striatum. J Neurochem. 2008;107:544–556.

